# Defining the spectrum of cellular protection against cadmium stress by Dsup expressed in yeast

**DOI:** 10.64898/2026.09.28.755128

**Authors:** Lewis Hitchens, Hamid Kian Gaikani, Marjan Barazandeh, Guri Giaever, Corey Nislow

## Abstract

The intrinsically disordered tardigrade Dsup protein has garnered attention for its ability to protect cells and organisms from radiation and oxidative stress. Our lab has previously described Dsup’s protective effect in yeast, finding that, while Dsup expression comes with a fitness cost, it improves survival under many stressors, including the heavy metal cadmium. While Dsup’s protection against oxidative stress and radiation is well-established, the mechanism by which it protects against other stressors is underinvestigated. To address this knowledge gap, we screened a collection of 4000 yeast deletion mutants expressing Dsup in the presence of three doses of cadmium to identify synergistic/synthetically lethal genes and processes. We find that Dsup broadly buffers many processes/genes at lower levels of cadmium stress but at higher doses, cells shift to buffering specific genes/processes. These results identify host genetic processes that influence Dsup’s ability to confer cadmium resistance to organisms.

**Article Summary:** Damage suppressor protein (Dsup) from extremophilic tardigrades protects against DNA damage when expressed, but how it protects against toxic metal stress is unclear. We screened 4,000 yeast deletion strains expressing Dsup at three doses of cadmium, a toxic heavy metal with deleterious biological and ecological effects. At low doses, Dsup buffered the loss of genes and processes while at higher doses buffering was restricted to specific genes. This shows genome-wide response to cadmium toxicity in yeast and identifies genes that interact with Dsup to confer protection. These observations highlight key affected processes and may inform efforts to engineer countermeasures.

## Introduction

Tardigrades, also known as water bears or moss piglets, represent a phylum of microscopic animals found ubiquitously in aquatic and terrestrial environments (Møbjerg et al. 2011). Some tardigrade species are highly extremotolerant, able to survive high heat, desiccation, and even the radiation of space (Møbjerg et al., 2011; Hashimoto and Kunieda, 2017). Dsup (Damage Suppressor protein) was initially characterized in *Ramazzottius varieornatus*, an extremophilic tardigrade species (Hashimoto et al., 2016). In that work, heterologous expression of Dsup in cultured mammalian cells was able to protect their DNA from high doses of ionizing radiation. Numerous studies have since assessed Dsup’s potential to protect cells and organisms from radiation and oxidative stress (Chavez et al., 2019; Kirke et al., 2020; Aguilar et al., 2025; Richaud et al., 2025). While there are species-specific variations in the extent of protection, the consensus is that Dsup acts by binding to DNA (or chromatin) and can confer stress resistance to mammalian cells, yeast, plants, and nematodes. Recently, researchers used lipid nanoparticle-encapsulated Dsup mRNA to deliver Dsup to mice with oral cancer (Kirtane et al., 2025). This preliminary study showed the potential biomedical applications of Dsup and suggests that the treatment can protect healthy tissues from radiation damage while preserving the tumor-killing efficacy of the treatment.

Dsup has many exciting potential applications in healthcare or for engineering stress resistant organisms, yet the mechanisms of its protective effect are still not fully understood. Furthermore, in those cases where Dsup expression appears deleterious, more work is needed. For example, one study found that exogenous Dsup causes neurotoxicity in primary cortical rat neurons (Escarcega et al., 2023). Our lab has extensively phenotyped Dsup in budding yeast, finding that Dsup can provide protection against an array of DNA-damaging agents, but that expression of Dsup comes at a fitness cost (Gaikani et al., 2025). We tested Dsup’s protective effect with a host of DNA-damaging agents including cadmium, a heavy metal pollutant (Järup and Åkesson, 2009; Zhang and Reynolds, 2019), and found a strong protective effect of Dsup against cadmium-induced stress. Cadmium is a ubiquitous environmental contaminant and carcinogen that can bioaccumulate in plants and animals (Zhang and Reynolds, 2019; Genchi et al., 2020). In humans cadmium can accumulate and cause systemic health issues (particularly in the liver and kidneys), mainly through consumption of food sourced from soils or environments with cadmium contamination (Järup and Åkesson, 2009; Zhang and Reynolds, 2019; Genchi et al., 2020). Prior to this work, Dsup’s protective effect had only been tested with heavy metal stress using arsenite, which has a similar but distinct mechanism of toxicity to cadmium (Thorsen et al., 2009; Đukić-Ćosić et al., 2020; Ozturk et al., 2021; Richaud et al., 2025). As such, we designed a chemogenomic screen to elucidate on the mechanisms and processes involved in this protective phenotype (Giaever and Nislow, 2014; Barazandeh et al., 2025). We screened a yeast knockout collection with an integrated Dsup-GFP (Dsup YKO) construct under the control of a β-estradiol-inducible promoter with three concentrations of cadmium chloride (at an approximately IC15, IC25, and IC35 respectively) (Sanford et al., 2022). In parallel, a knockout collection with GFP (GFP YKO) under the control of the same β-estradiol-inducible promoter was screened as a control.

## Methods

### Preparing the Assay

We used the Dsup YKO and GFP YKO generated from our previous work via synthetic genetic array (SGA)(Gaikani et al., 2025). An aliquot of the Dsup YKO collection and one of the GFP YKO collection was thawed from –70 °C and diluted in SC media to an OD_600_ of 0.06 (from a concentrated stock at an OD_600_ of 40). Each YKO culture was split into two portions: (0 nM β-estradiol induction and 40 nM β-estradiol induction). 40 nM was chosen for Dsup induction based on our lab’s previous Dsup YKO bar-seq assay. We adapted standard bar-seq/chemogenomics assays: 686 μl of cell culture was added to wells in a 48-well plate along with 14 μl of 50x cadmium chloride solution (Suresh et al., 2016; Barazandeh et al., 2025). For untreated controls, 14 μl of water was added instead of cadmium. Each plate was covered with a clear seal (Thermo Scientific, AB0580) before being placed into a GP2 custom liquid handling robot (S&P Robotics, Toronto, ON, Canada). First, a bioassay was performed with the Dsup YKO to determine dose response of cadmium with a serial halving dilution of cadmium chloride (7 concentrations ranging from 7.8125-500 μM), and growth curves were analyzed using the AUDIT platform (https://ggshiny.shinyapps.io/AUDIT-geneticnetworks/)(Coutin et al., 2020). Concentrations of cadmium used for the subsequent bar-seq assay (each condition in triplicate) were 15.625 μM, 31.25 μM, and 62.5 μM (rounded to one decimal place for clarity in all figures). These concentrations corresponded to approximately 15%, 25%, and 35% inhibition respectively (Figure S1).

### GP2 Homozygous Profiling (HOP) Assay

The GP2 maintained cells at 30 °C with intermittent shaking. Cells were grown for ∼5 generations (until OD_600_ of 1.2), and then collected by the robot and stored at 4 °C. OD_600_ was monitored every 10-15 minutes. After all samples were collected, genomic DNA was extracted using the Yeastar Genomic DNA kit (Zymo Research, D2002).

### PCR Reactions

Strain-specific barcodes were amplified from extracted gDNA and indexed with

Illumina-compatible adapters in two PCR reactions sequentially. PCR1 amplified both the uptags and downtags which were then pooled together for PCR2 which added the Illumina adapters. Each PCR reaction had a total volume of 20 µL, containing 0.2 µL of Thermo Scientific Phusion Hot Start II DNA polymerase (2 U/µL), 4 µL of 5× Phusion HF buffer, 0.4 µL of 10 mM dNTPs, 80 ng of genomic DNA, and 0.5 µM of forward and reverse primers. Thermocycling conditions: initial denaturation: 3 min at 98 °C, 25 cycles of denaturation at 98 °C for 10 s, annealing at 59 °C for 30 s, extension at 72 °C for 20 s, and final extension at 72 °C for 5 min. PCR2 reactions were performed in a total volume of 50 µL, containing 0.5 µL of Phusion Hot Start II DNA polymerase, 10 µL of 5× Phusion HF buffer, 1 µL of 10 mM dNTPs, 1.5 µL of 1/10 diluted PCR1 product, and 0.5 µM of PCR2 primers. For PCR2, thermocycling conditions were as follows: 3 min at 98 °C, 6 cycles of denaturation at 98 °C for 30 s, annealing at 60 °C for 30 s and extension at 72 °C for 30 s, and final extension at 72 °C for 5 min.

### PCR1 Illumina uptag primers

5′ TCGTCGGCAGCGTCAGATGTGTATAAGAGACAGGATGTCCACGAGGTCTCT 3′

5′ GTCTCGTGGGCTCGGAGATGTGTATAAGAGACAGGTCGACCTGCAGCGTACG 3′

### PCR1 Illumina downtag primers

5′ TCGTCGGCAGCGTCAGATGTGTATAAGAGACAGGAAAACGAGCTCGAATTCATCG

3′

5′ GTCTCGTGGGCTCGGAGATGTGTATAAGAGACAGCGGTGTCGGTCTCGTAG 3′

### PCR2 primers for Illumina sequencing

5′ CAAGCAGAAGACGGCATACGAGAT[i7]GTCTCGTGGGCTCGG 3′

5′ AATGATACGGCGACCACCGAGATCTACAC[i5]TCGTCGGCAGCGTC 3

### Sequencing and Statistics

Libraries were pooled, normalized, and sequenced on the NovaSeq X Plus platform (Paired End 150), targeting 5 to 6 million reads per sample. Analysis was performed with the pipeline our lab described for bar-seq (Barazandeh et al., 2025). For statistical comparison of Normalized Enrichment Score (NES) magnitude between Dsup-/Dsup+ and GFP-/GFP+ cells respectively, processes were filtered for False Discovery Rate (FDR) q-value < 0.05 in at least one of the two conditions for each dose. Analyses were performed and visualized in R v4.5.2 with various packages (dplyr, ggplot2, patchwork, tidyr) (Wickham, François, et al., 2014; Wickham, Vaughan, et al., 2014; Wickham, 2016; Pedersen, 2019).

### Gene Set Enrichment Analysis

Fitness defect (FD) scores were calculated by normalizing log₂ ratios of tag counts between drug-treated samples and controls, with positive values indicating sensitivity and negative values indicating resistance to the treatment. Pre-ranked GSEA was performed with GSEA v4.4.0 using a minimum gene set size of 10 and maximum size of 300 using the *S. cerevisiae* GO biological processes file from UBC EasyGSEA (Subramanian et al., 2005; Cheng et al., 2021).

## Results and Discussion

### Dsup acts as a broad genetic buffer under mild but not high levels of cadmium stress

Our previous work showed that Dsup can provide protection from a variety of stressors, including cadmium and hydrogen peroxide-induced oxidative stress (Gaikani et al., 2025). While cadmium exposure leads to oxidative stress, it also directly causes proteotoxic stress by binding to sulfhydryl groups and may displace other divalent cations from binding sites, disrupting protein function (Wysocki and Tamás, 2010; Tamás et al., 2014; Ozturk et al., 2021). As such, the global effects of cadmium stress require a large number of processes to be functional to adequately defend against its toxicity (Vido et al., 2001; Ruotolo et al., 2008; Thorsen et al., 2009; Wysocki and Tamás, 2010). We looked at how the median of normalized enrichment score (NES) for GO biological processes (with FDR q-value < 0.05) changes from Dsup-to Dsup+ and across doses to identify trends in global enrichment (Figure 2). We found that moderate Dsup expression can provide broad genetic buffering across processes at the lowest dose of cadmium tested (i.e., a dose that causes ∼15% fitness defect), but this buffering effect declines as cadmium concentration increases. (Figures 2a, 2b). At 31.3 μM cadmium chloride (approx. 25% inhibition), Dsup can buffer median global enrichment increases (i.e., processes imposing a fitness defect) with cadmium dose but does not significantly change negative enrichment (i.e., processes imposing a fitness advantage). In other words, at low doses Dsup-expressing cells manifest a decrease in enrichment magnitude of affected GO terms, but they do not at higher doses. When the control collection of GFP-expressing yeast deletion mutants was subjected to cadmium stress, GFP+ cells had an increased median NES for biological processes with FDR q-value < 0.05 at a low cadmium concentration and high cadmium concentration, suggesting that our findings with Dsup are Dsup-specific and not simply the result of heterologous expression of any protein causing shifts in process dynamics. These observations are consistent with a model where Dsup can globally buffer cellular fitness at low doses of a cadmium but shifts to more specific processes/mechanisms to improve cellular survival (as opposed to fitness) at higher doses. For example, the broad buffering observed at lower doses shifts to buffering of key individual genes in the cadmium response like *GSH2* and *YCF1* at higher doses (Figures 2a, 4). GFP expression had the opposite effect, pushing median enrichment higher than the uninduced control (at the lowest dose) (Figures 2a, 2c), which further supports the model of Dsup as a broad buffer.

**Figure 1.**
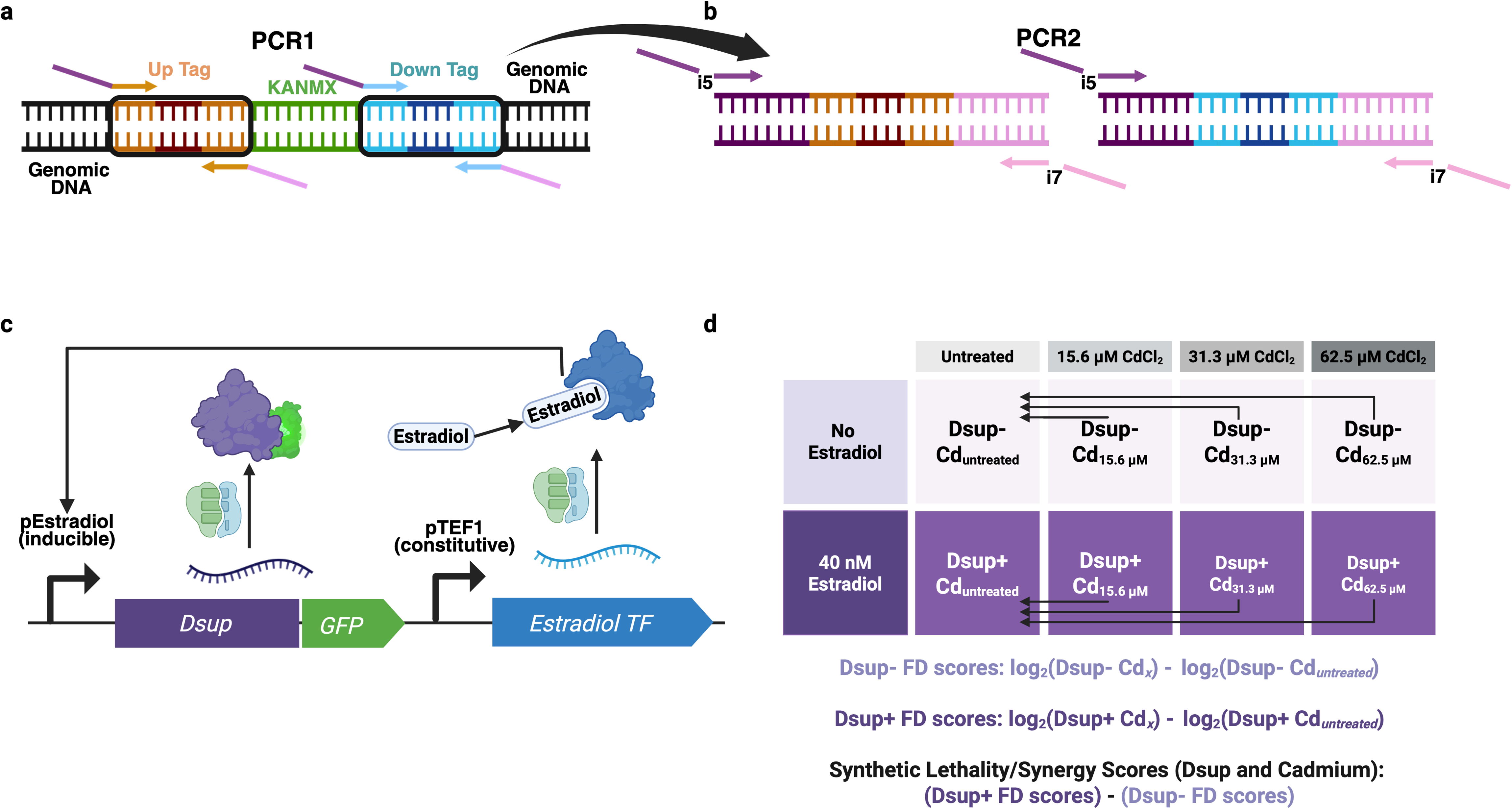
Overview of experimental procedure. a) PCR1 amplifies uptags and downtags at the locus of each respective gene deletion (in separate PCR reactions). b) PCR2 takes the pooled uptags and downtags and adds Illumina indices for sequencing. c) The β-estradiol inducible system for expressing Dsup, integrated at the (deleted) *LEU2* locus. A transcription factor (from plasmid pAS161, Addgene #194484) with a β-estradiol-binding domain is transcribed constitutively. When β-estradiol binds the transcription factor a chaperone dissociates, allowing the transcription factor to enter the nucleus where it binds to response elements in pEstradiol (from plasmid pAS165, Addgene #194488), initiating transcription of our downstream Dsup-GFP construct. pAS161 and pAS165 are domesticated into the Yeast MoClo Toolkit which was employed to clone the plasmids used to integrate the codon-optimized Dsup vector for construction of the Dsup YKO (Lee et al. 2015; Sanford et al. 2022; Gaikani et al. 2025). d) Conditions in the Dsup bar-seq screen and comparisons made to generate fitness defect (FD) scores from normalized log_2_ ratios of tag counts, which were then used to calculate synthetic lethality and synergy scores. Panels a and b adapted from Barazandeh et al. (2025). Created in BioRender. Hitchens, L. (2026) https://BioRender.com/2n2sc0t **Alt Text:** Graphic illustration of experimental procedure, with panel a showing how uptags and downtags are amplified for PCR1 at each deletion locus using a visual of double stranded DNA. Panel b shows how Illumina adapters are added to each tag with a visual of double stranded DNA in the same style. Panel c shows the β-estradiol-inducible Dsup-GFP construct with Dsup in purple, GFP in green, and the β-estradiol-binding transcription factor in blue. Arrows and icons denoting transcripts, ribosomes, and proteins illustrate how the system works to drive expression of the Dsup construct. Panel d shows the different conditions as a labelled 2×4 matrix in the Dsup bar-seq screen with equations below used to calculate fitness defect scores which were then used to calculate synthetic lethality and synergy scores.

**Figure 2.**
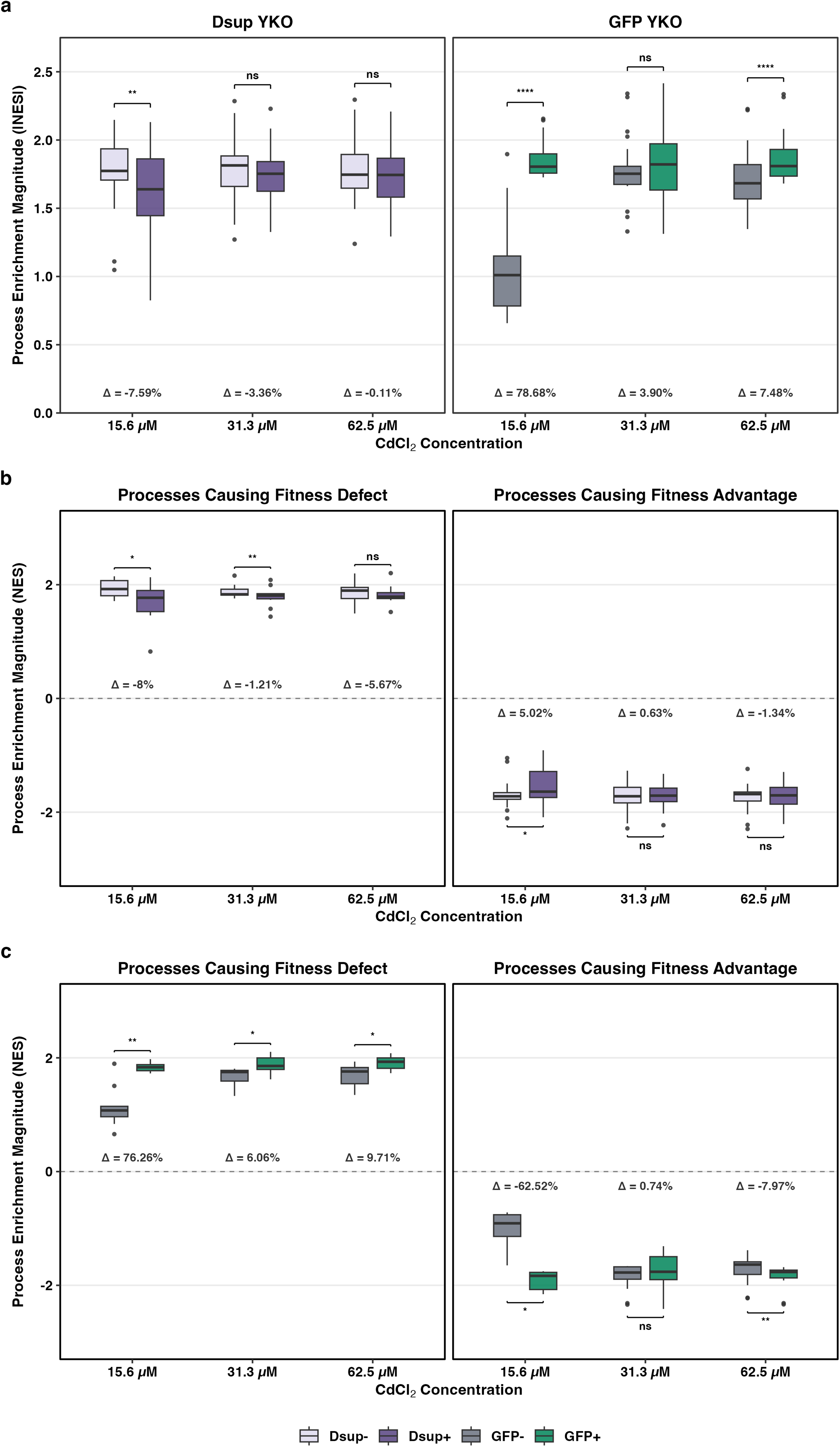
Dsup’s global genetic buffering across cadmium doses. a) Comparison of process enrichment magnitudes (processes only included if FDR q-value < 0.05 in at least one condition, i.e. Dsup-/Dsup+ or GFP-/GFP+) from pre-ranked GSEA analyses comparing cadmium exposed Dsup YKO and GFP YKO cells to their respective controls (e.g., Dsup+Cd+ vs. Dsup−Cd+ and Dsup+Cd− vs. Dsup−Cd−). Boxplots represent interquartile range (IQR) and median (center line). Dsup- are cells with no β-estradiol induction of Dsup. Dsup+ are cells with 40 nM β-estradiol induction of Dsup. Statistical comparisons performed with two-sided Wilcoxon signed-rank test. * = p < 0.05; ** = p < 0.01; *** = p<0.001; **** = p<0.0001; ns = not significant. NES = Normalized Enrichment Score. Δ = percent change in median |NES| from Dsup- to Dsup+ or GFP- to GFP+ respectively. Dots represent outlier processes. b) Comparison of direction-dependent enrichment magnitudes for the Dsup YKO, grouped as paired sets based on initial sign (i.e., negative or positive) at baseline in Dsup-. Boxplots represent IQR and median (center line). Statistical comparisons were performed with a two-sided Wilcoxon signed-rank test. NES = Normalized Enrichment Score. * = p < 0.05; ** = p < 0.01; *** = p < 0.001; **** = p < 0.0001; ns = not significant. NES = Normalized Enrichment Score. Δ = percent change in median NES from Dsup- to Dsup+. Dots represent outlier processes. c) Comparison of direction-dependent enrichment magnitudes for the GFP YKO. Boxplots represent IQR and median (center line). To avoid distortion by processes that sign-flip from GFP- to GFP+, processes are strictly directionally bounded. Statistical comparisons performed with a two-sided Wilcoxon signed-rank test. NES = Normalized Enrichment Score. * = p < 0.05; ** = p < 0.01; *** = p < 0.001; **** = p < 0.0001; ns = not significant. NES = Normalized Enrichment Score. Δ = percent change in median NES from GFP- to GFP+. Dots represent outlier processes. **Alt Text:** Three-panelled figure of two-faceted boxplots depicting magnitude of enrichment for GO biological processes. Light purple bars represent Dsup-, darker purple represents Dsup+, grey represents GFP-, and green represents GFP+ Panel a depicts absolute value of NES with the Dsup YKO on the left facet, and the GFP YKO cells on the right facet. Panel b shows direction-dependent magnitudes of enrichment for Dsup YKO cells with positive enrichment on the left facet, and negative enrichment on the right facet. Panel c shows direction dependent magnitudes of enrichment for GFP YKO cells with positive enrichment on the left facet, and negative enrichment on the right facet.

### Dsup can partially buffer some vacuolar and protein targeting deficiencies under mild cadmium stress

A primary cellular detoxification mechanism for cadmium is sequestration of the metal to the vacuole (Mendoza-Cózatl et al., 2005; Wysocki and Tamás, 2010; Ozturk et al., 2021). We found that at a low dose of cadmium, Dsup is able to buffer two protein targeting and vacuolar processes: “Vacuolar Acidification”, and “Retrograde Transport, Endosome to Golgi” (Figure 3). Deletion of genes in these processes confers a relative fitness advantage when Dsup is expressed under 15.6 μM of cadmium exposure. Specifically, genes like *PEP8, YPT6, RAV1*, and *MEH1* are directly or indirectly involved in heavy metal tolerance (Table 1) (Ruotolo et al., 2008; Zhang et al., 2017). Vacuolar acidification via V-ATPase proton pumps helps balance redox under metal stress to maintain the activity of some antioxidant proteins (Techo et al., 2020). Dsup could be mitigating oxidative stress from cadmium to partially buffer this process.

**Figure 3.**
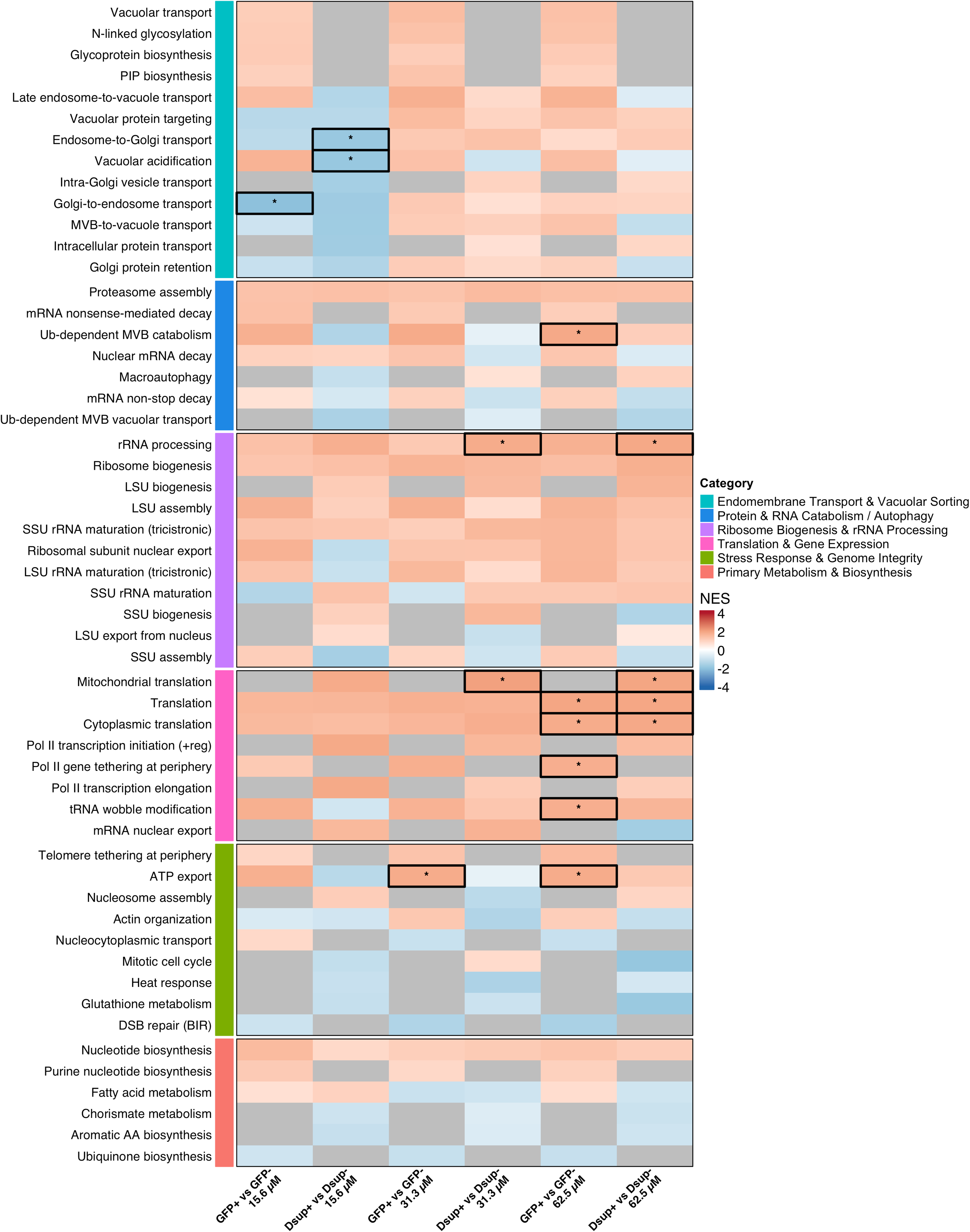
Heatmap of Enriched Processes. Enriched processes (filtered for FDR q < 0.05 in at least 1 out of 18 respective GFP or Dsup conditions) for synthetic lethality scores made for GFP YKO and Dsup YKO. Blue represents a relative fitness advantage, and red represents a relative defect. Grey represents processes that were not given an enrichment score by GSEA for a given condition due to insufficient representation (processes with insufficient representation for the six visualized conditions were removed). Asterisks represent processes with FDR q < 0.05 for each respective condition. Processes are from GO biological processes for *S. cerevisiae* from UBC EasyGSEA. Some longer GO biological processes were abbreviated as needed for visualization, with full length names and GO IDs available in Supplementary Table 1. **Alt Text:** Heatmap of GO biological processes for Dsup and GFP synthetic lethality scores grouped into broad categories by colored bars on the left, depicted as a six-column matrix. Positive enrichment is represented as red, and negative enrichment is represented as blue, with the intensity of each color corresponding to magnitude of enrichment. Asterisks and black boxes highlight biological pathways with FDR q < 0.05.

**Table 1.** Genes (FD threshold +/- 1) for each process with synthetic lethality/synergy (with FDR q < 0.05) for Dsup. If a process that was enriched with q > 0.05 that did not have genes with fitness scores less than −1 or greater than 1, they were assigned as N/A in the “Gene” and “FD Score” column. Note: For genes in Retrograde Transport, Endosome to Golgi, *YPT6* had an FD score of –0.98; for Vacuolar Acidification, top genes were *RAV1* and *MEH1* both with an FD score of –0.68. Shades of purple for processes are distinct solely for visual clarity. Shades of red represent the intensity of synthetic lethality. All scores were rounded to two decimal places.

| Condition | Biological Process | Gene | FD Score |
| --- | --- | --- | --- |
| Dsup 15.6 μM | Retrograde Transport, Endosome to Golgi | *PEP8* | -1.14 |
| Dsup 15.6 μM | Vacuolar Acidification | N/A | N/A |
| Dsup 31.3 μM | Mitochondrial Translation | *MRX14* | 1.77 |
| Dsup 31.3 μM | rRNA Processing | *FYV7* | 3.13 |
| Dsup 31.3 μM | rRNA Processing | *RPL8B* | 1.85 |
| Dsup 31.3 μM | rRNA Processing | *BUD27* | 1.22 |
| Dsup 31.3 μM | rRNA Processing | *RPL8A* | 1.09 |
| Dsup 62.5 μM | Cytoplasmic Translation | *RPL8B* | 2.30 |
| Dsup 62.5 μM | Cytoplasmic Translation | *RPL22A* | 1.75 |
| Dsup 62.5 μM | Cytoplasmic Translation | *RPS7A* | 1.55 |
| Dsup 62.5 μM | Cytoplasmic Translation | *RPS10A* | 1.45 |
| Dsup 62.5 μM | Cytoplasmic Translation | *RPS28B* | 1.28 |
| Dsup 62.5 μM | Cytoplasmic Translation | *RPS25A* | 1.22 |
| Dsup 62.5 μM | Cytoplasmic Translation | *RPL19B* | 1.04 |
| Dsup 62.5 μM | Cytoplasmic Translation | *RPL33B* | 1.00 |
| Dsup 62.5 μM | Mitochondrial Translation | *MRX14* | 2.70 |
| Dsup 62.5 μM | Mitochondrial Translation | *RSM18* | 1.46 |
| Dsup 62.5 μM | Mitochondrial Translation | *SLS1* | 1.37 |
| Dsup 62.5 μM | Mitochondrial Translation | *RSM27* | 1.30 |
| Dsup 62.5 μM | rRNA Processing | *FYV7* | 3.30 |
| Dsup 62.5 μM | rRNA Processing | *RPL8B* | 2.30 |
| Dsup 62.5 μM | rRNA Processing | *BUD27* | 1.72 |
| Dsup 62.5 μM | rRNA Processing | *RPS7A* | 1.55 |
| Dsup 62.5 μM | rRNA Processing | *NSR1* | 1.32 |
| Dsup 62.5 μM | rRNA Processing | *DBP3* | 1.00 |
| Dsup 62.5 μM | Translation | *MRX14* | 2.70 |
| Dsup 62.5 μM | Translation | *RPL22A* | 1.76 |
| Dsup 62.5 μM | Translation | *RPS7A* | 1.55 |
| Dsup 62.5 μM | Translation | *RSM18* | 1.46 |
| Dsup 62.5 μM | Translation | *RPS28B* | 1.28 |
| Dsup 62.5 μM | Translation | *RPL19B* | 1.04 |
| Dsup 62.5 μM | Translation | *RPL33B* | 1.00 |

Endosome-to-Golgi transport is responsible for recycling vacuolar trafficking components and perturbations in this process may impair metal detoxification by causing protein missorting, impairing vacuole functionality (Ruotolo et al., 2008; Ma and Burd, 2020). This may reflect Dsup’s protective effect which limits the reliance on these processes for detoxification of cadmium by mitigating the oxidative stress caused by cadmium exposure, and therefore compensating for the lack of intact vacuolar and trafficking processes to detoxify cadmium.

### Dsup and GFP expression abrogates cadmium resistance in translation-deficient mutants at higher cadmium concentrations, suggesting that Dsup’s protection against cadmium imposes a similar protein burden as seen with a “neutral” protein

At higher concentrations of cadmium (31.3 μM, 62.5 μM), we observed that deletion of genes involved in both cytoplasmic and mitochondrial translation manifested fitness defects (Figure 3). This observation is consistent with cadmium targeting the mitochondria by binding to the thiol groups on complexes, uncoupling the Electron Transport Chain (ETC) which, in turn, leads to ROS generation that causes the oxidative stress associated with cadmium treatment (Branca et al., 2020). Impaired mitochondrial translation accuracy can increase accumulation of ROS and protein aggregation in yeast (Suhm et al., 2018). Our previous research has shown that Dsup is localized exclusively to the nucleus and not the mitochondria in *S. cerevisiae*, so it is unlikely that mitochondrial translation processes are directly interacting with Dsup (Gaikani et al., 2025). While Dsup expression imposes a fitness cost for deletions in some translation genes and/or translation-adjacent processes, similar translation-related processes are positively enriched (i.e., imposing a relative fitness defect) under cadmium stress in the Dsup YKO compared to the GFP YKO. It is important to note that even though GFP may be considered an “inert” or “neutral” protein, heterologous expression of GFP and other fluorescent proteins have been shown to cause protein burden when overexpressed (Fujita et al., 2025). Specifically, we found that expressing GFP imposes a similar reliance on translational and ribosomal machinery compared with Dsup expression under cadmium stress. Therefore, heterologous Dsup expression does not impose a notably higher burden on translational or ribosomal machinery under cadmium stress than a generic/neutral protein in GFP.

### Dsup expression can reduce reliance on key genes involved in cadmium detoxification

While we did not find processes conferring a relative fitness advantage/synergy to be enriched in the 31.3 and 62.5 μM cadmium doses, several key genes showed a relative fitness advantage when Dsup was expressed in these higher cadmium doses (Figure 4). *GSH2* is glutathione synthetase which catalyzes the synthesis of glutathione, the primary chelator for cadmium and a key antioxidant enzyme for defense against oxidative stress (Inoue et al., 1998; Vido et al., 2001; Dai et al., 2024). *YCF1* is yeast cadmium factor 1, which transports the glutathione-cadmium conjugate into the vacuole (Mendoza-Cózatl et al., 2005; Dai et al., 2024). *IRE1* mediates the unfolded protein response that occurs under cadmium exposure (Gardarin et al., 2010; Le et al., 2016). Some previously underexplored genes are also shown to provide a relative fitness advantage when knocked out under cadmium stress. *VPS61* is a dubious ORF directly upstream of *YCF1* whose deletion is known to cause vacuolar protein sorting defects (Bonangelino et al., 2002). *YML034C-A* is another dubious reading frame that shows this phenotype but has an unknown function. Combined with the deterioration of the global buffering effect under increased cadmium dose, this suggests a system where Dsup’s broad protection at low cadmium doses transitions to highly process/gene-specific buffering to aid in survival at higher doses.

**Figure 4.**
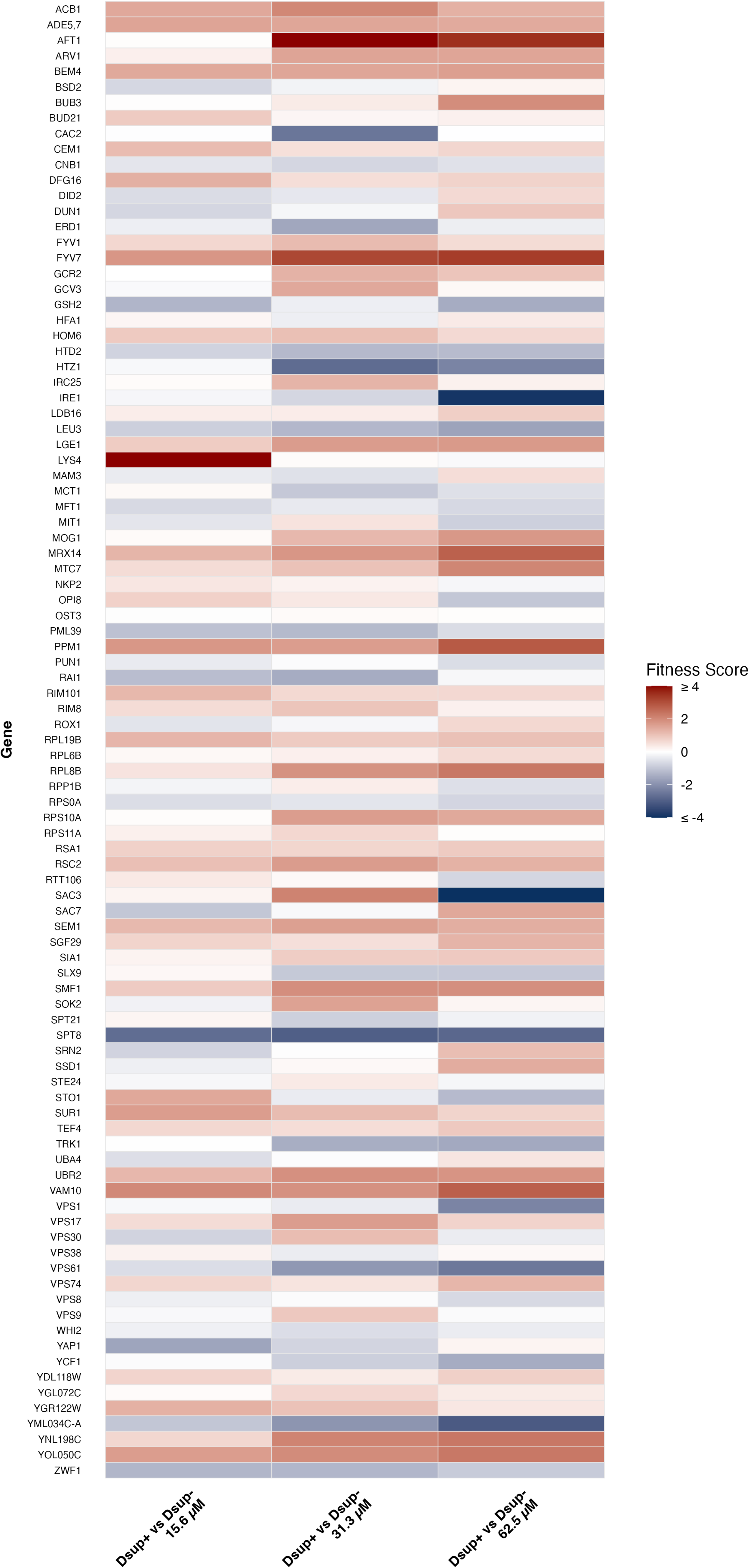
Heatmap of Dsup synthetic lethality/synergy scores for individual genes. Red represents positive enrichment (i.e., a relative) fitness defect/synthetic lethality. Blue represents negative enrichment (i.e., a relative) fitness advantage/synergy. Genes were filtered for FD ≥ 2 or ≤ −2 in at least one of the 9 conditions (i.e., Dsup-, Dsup+, or Dsup synthetic lethality scores for each cadmium dose). **Alt Text:** Heatmap of synthetically lethal or synergistic genes for Dsup YKO for all three cadmium doses, depicted as a three-column matrix. Positive enrichment is represented as red, and negative enrichment is represented as blue, with the intensity of each color corresponding to magnitude of enrichment.

## Conclusion

These findings suggest that Dsup’s protective effect against cadmium stress in a model eukaryotic organism is dose-dependent and shifts from providing broad fitness buffering at low cadmium doses to gene-specific buffering at higher doses of cadmium. We have also shown that while Dsup’s protective effect under higher levels of cadmium stress does rely on an intact translational machinery, its expression imposes a similar burden on translational processes to a “neutral” protein in GFP.

Our lab has previously explored the trade-off of fitness for survival that Dsup imposes in eukaryotic systems across several DNA-damaging drugs (Gaikani et al., 2025). By understanding the processes involved/affected by Dsup’s mechanism of action across multiple doses of cadmium, we provide a genome-wide profile of how this previously determined trade-off manifests under increasing levels of exposure to a previously uninvestigated stressor.

The scope of this study is limited to fitness of knockouts under cadmium stress rather than survival of Dsup-expressing strains (which was investigated in our previous work Gaikani et al., 2025). Whether Dsup is acting as a mitigator of oxidative stress to improve fitness under cadmium stress or is protecting genome integrity from cadmium in another manner remains to be investigated. It will also be important to investigate whether Dsup can improve uptake of cadmium or other heavy metals.

Given our lab’s findings, together with the previous investigation of arsenite exposure in Dsup-expressing nematodes, it stands to reason that Dsup expression may be able to protect organisms from other heavy metals like mercury, lead, or chromium (Gaikani et al., 2025; Richaud et al., 2025). Previous work on Dsup has largely focused on protection from radiation and oxidative stress, but this work underscores Dsup’s potential for use in engineering organisms for bioremediation of pollutants. Heterologous expression of a tapeworm metallothionein has previously been shown to boost accumulation of cadmium in *E. coli* (He et al., 2024).

Additionally, *S. cerevisiae* has been engineered to hyperaccumulate cadmium by 1-2 orders of magnitude (Sun et al., 2019), and Dsup expression might be able to act as a multiplier for yeast or other microbial hyperaccumulators. In conjunction with our previous work, these findings provide a systemic genetic picture of Dsup’s potential for use in protecting cells and organisms from cadmium.

## Data Availability Statement

All sequencing data associated with this study have been deposited in the NCBI Sequence Read Archive (SRA) as BioProject PRJNA1534049 (accessions for each sample: SAMN63429291– SAMN63429338).

## Supplementary Data

**Figure S1. Bioassay results for Dsup YKO.** Growth inhibition results (using Dsup-untreated cells as references) for Dsup YKO when tested with a range of cadmium chloride concentrations, Dsup+ represents 75 nM of β-estradiol induction. In our previous work Gaikani et al. (2025), we used 75 nM β-estradiol induction for experiments measuring survival under stress but 40 nM induction for bar-seq screens to confer a more modest Dsup-associated fitness defect and increase dynamic range of the screen. Y-axis measures OD_600_ as measured from the plate reader in GP2 robot, x-axis measures runtime in hours. Initial visualization generated with AUDIT platform (https://ggshiny.shinyapps.io/AUDIT-geneticnetworks/). Partially created in BioRender. Hitchens, L. (2026) https://BioRender.com/oz6a5hx

**Figure S1 Alt Text:** Inhibition curves and percentages for Dsup YKO (using Dsup-untreated cells as references) under cadmium stress as a 6×8 matrix, light purple box outlines Dsup-cells, and a darker purple box outlines Dsup+ cells (75 nM β-estradiol induction). Cadmium doses increasing from left to right are labelled at the top of the figure.

---

**Supplementary Table 1.**
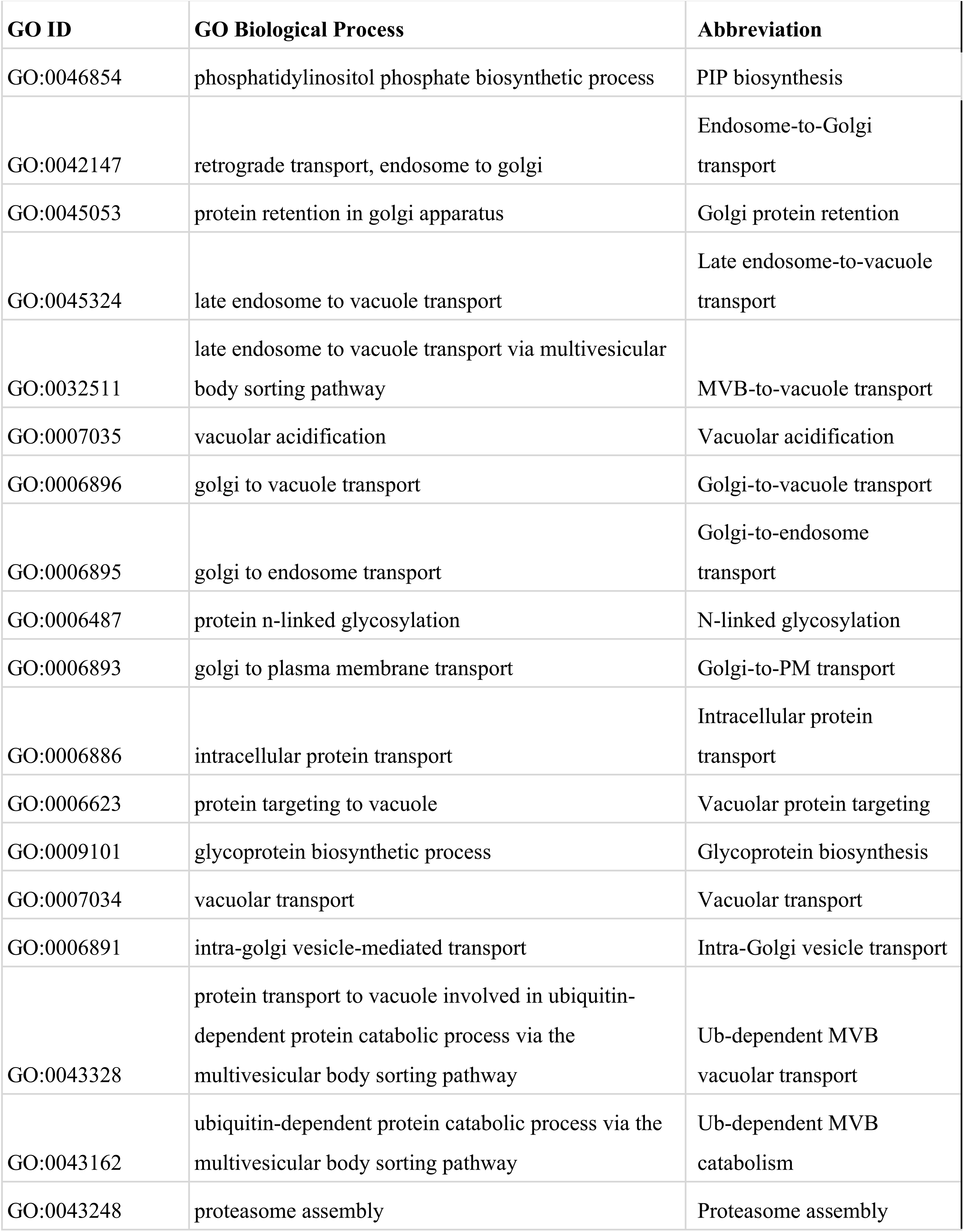
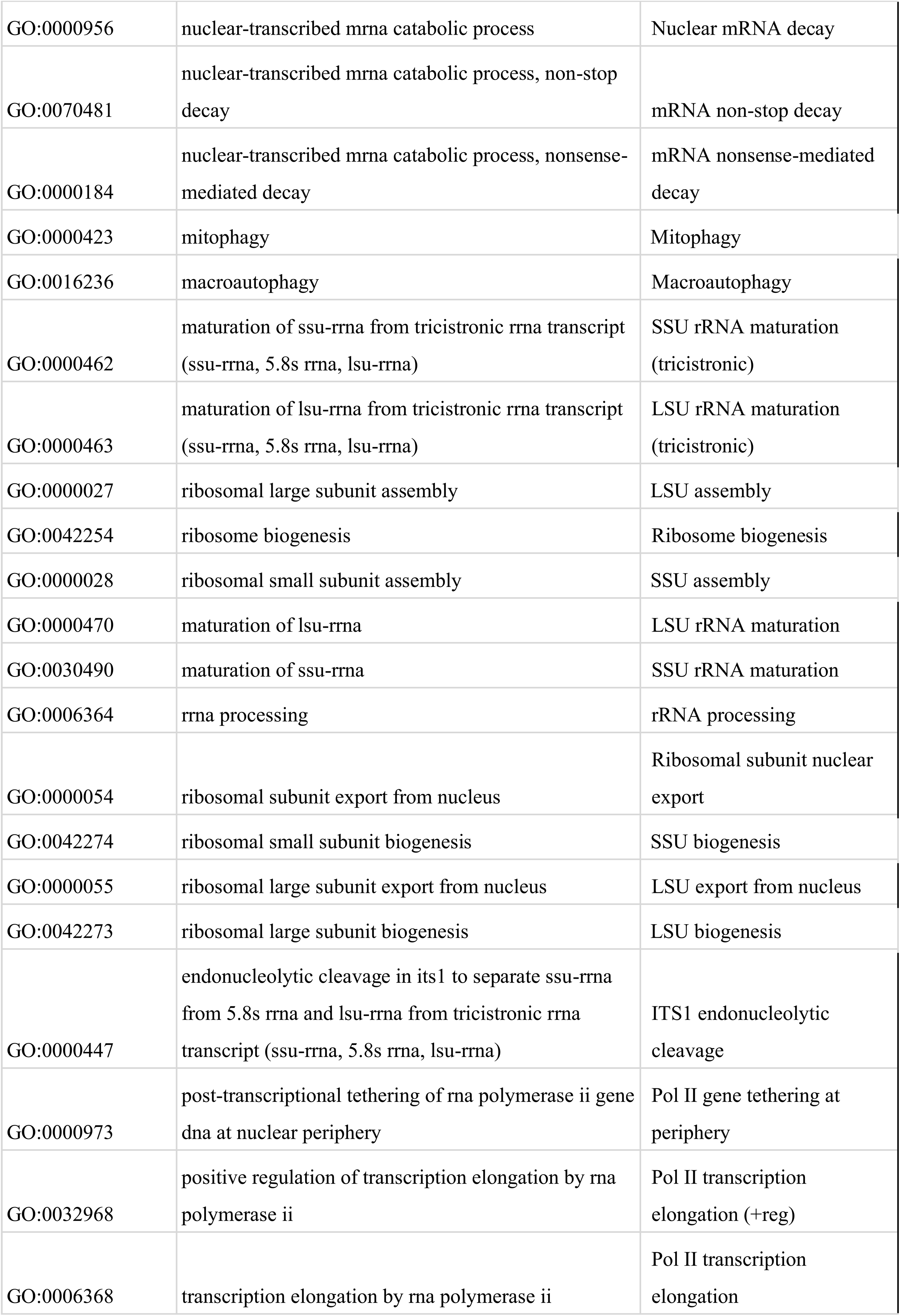
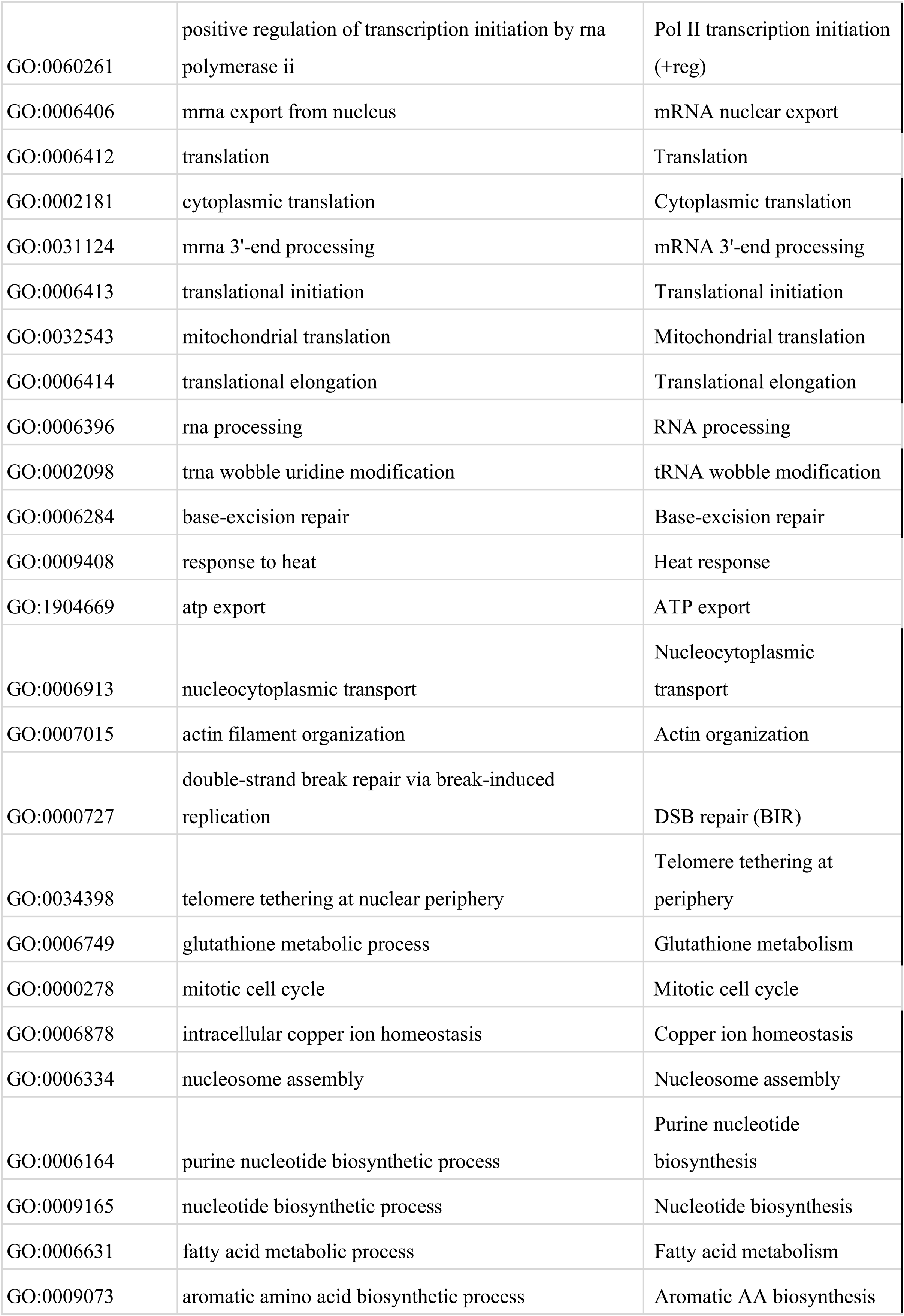
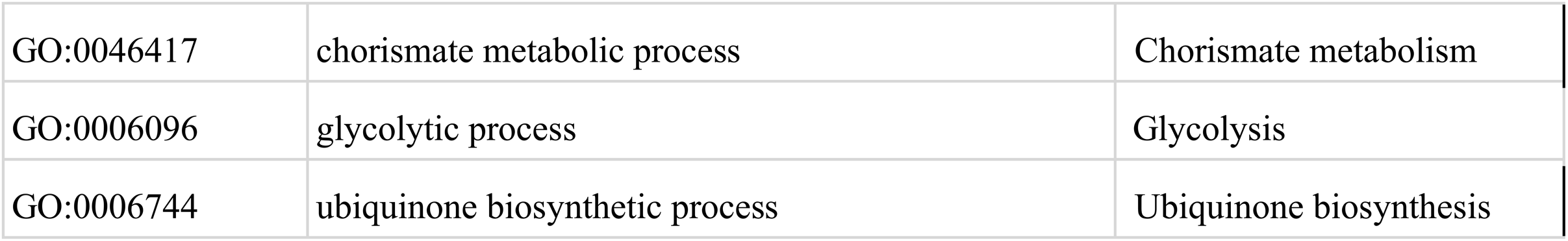
GO Terms and Abbreviations. Abbreviations of GO Biological processes used in Figure 3 alongside full-length names and GO IDs. Note that this table also includes processes with insufficient representation for all six visualized conditions in Figure 3 that were represented in other conditions with FDR q < 0 in Figure 3 that were represented in other conditions with FDR q < 0.05.

